# Voluntary wheel running selectively preserves skeletal muscle during pancreatic cancer cachexia despite persistent systemic alterations

**DOI:** 10.64898/2026.10.14.758481

**Authors:** Cindy Richard, Arthur Dieumegard, Lou-Anne Cavrois, Luz Lefeuvre-Orfila, Brice Martin, Laurent Rouvière, Stéphane Servais, Manuel Vlach, Gaëlle Angenard, Catherine Ribault, Anne Corlu, Pascal Loyer, Amélie Rébillard

**Author notes:** Corresponding author: Prof. Amélie Rébillard, Laboratoire Mouvement, Sport, Santé (M2S), EA 7470 Campus de Ker Lann, Avenue Robert Schuman 35170 Bruz, FRANCE. These authors contributed equally to this work (Cindy Richard, Arthur Dieumegard).

## Abstract

**Key points:**

- Pancreatic cancer cachexia affects multiple physiological systems but whether these compartments retain similar responsiveness to exercise is unclear.
- Voluntary wheel running slowed tumor growth, reduced final tumor mass and preserved skeletal muscle mass whereas treadmill running did not reproduce these adaptations and was associated with a less favorable body-weight trajectory and greater fat mass loss.
- Skeletal muscle preservation with voluntary wheel running remained evident after adjustment for tumor burden indicating that reduced tumor burden alone was insufficient to account for the muscle response.
- In contrast, adipose tissue wasting was not prevented and was even accentuated at a given tumor burden, whereas hepatic metabolic alterations and systemic inflammation were largely unaffected by either exercise paradigm, revealing marked tissue-specific differences in physiological responsiveness.
- These findings indicate that skeletal muscle retains substantial exercise-induced plasticity during pancreatic cancer cachexia and that exercise responses depend on the characteristics of the exercise paradigm.

**Background:** Pancreatic ductal adenocarcinoma (PDAC) is frequently associated with cancer cachexia, involving alterations across multiple physiological systems. Whether distinct exercise paradigms similarly influence these compartments remains unclear. We compared voluntary wheel running (VWR) and treadmill running (TR) using an integrative physiological approach in a murine model of pancreatic cancer cachexia.

**Methods:** Male C57BL/6J mice bearing UN-KC-6141 pancreatic tumor cells were allocated to inactive control (CaCTL), VWR or TR groups and compared with healthy controls. Tumor progression, body composition, skeletal muscle, adipose tissue, hepatic metabolism, systemic inflammation, energy expenditure and behavior were assessed. Analysis of covariance determined whether tissue-mass differences remained evident after adjustment for tumor burden.

**Results:** UN-KC-6141 tumors induced a systemic cachectic phenotype involving skeletal muscle and adipose tissue wasting, hepatic metabolic remodeling, systemic inflammation and behavioral alterations. VWR slowed tumor growth, reduced final tumor mass, improved the body-weight trajectory and preserved lean and skeletal muscle mass, whereas TR did not reproduce these adaptations. Greater gastrocnemius mass with VWR remained evident after adjustment for tumor mass, indicating that reduced tumor burden alone was insufficient to account for muscle preservation. In contrast, adipose tissue wasting, hepatic metabolic remodeling and systemic inflammation remained largely unaffected. TR was also associated with a less favorable body-weight trajectory and greater fat mass loss than in inactive tumor-bearing mice. Exploratory metabolic phenotyping revealed distinct feeding, sleep and resting energy expenditure patterns between the two exercise paradigms.

**Conclusions:** VWR selectively preserved skeletal muscle within a persistently altered systemic environment, whereas the TR protocol used here did not reproduce these adaptations. These findings reveal tissue-specific exercise responsiveness during pancreatic cancer cachexia and highlight exercise paradigm characteristics as determinants of tumor-host adaptation.

## Introduction

Pancreatic ductal adenocarcinoma (PDAC) remains one of the deadliest human malignancies, with an overall five-year survival rate of approximately 13% despite substantial advances in diagnosis and treatment (American Cancer Society, 2026). Beyond its aggressive local invasion and metastatic dissemination, PDAC profoundly disrupts whole-body physiology and is among the malignancies most strongly associated with cancer cachexia (Baracos *et al*., 2018). Affecting up to 80% of patients with advanced disease (von Haehling *et al*., 2016), cachexia is a multifactorial syndrome defined by an ongoing loss of skeletal muscle mass, with or without loss of fat mass, that cannot be fully reversed by conventional nutritional support and leads to progressive functional impairment (Fearon *et al*., 2011). It impairs quality of life, reduces tolerance to anticancer therapies and is associated with increased mortality, yet effective strategies to preserve host physiology during pancreatic cancer progression remain limited.

Exercise has emerged as a non-pharmacological strategy to attenuate cancer-associated physical and metabolic dysfunction, but its efficacy against cachexia remains debated. A meta-analysis of preclinical studies found no significant overall effect of exercise on body or skeletal muscle mass, with substantial variability across tumor models and protocols (Niels *et al*., 2020). Nevertheless, recent studies indicate that skeletal muscle retains responsiveness to exercise during tumor progression, depending on the timing, duration and characteristics of exercise (Collao *et al*., 2023; Tsitkanou *et al*., 2026). Exercise should therefore not be considered a uniform intervention. Voluntary wheel running (VWR) and treadmill running (TR), two commonly used paradigms in rodents, differ in initiation, workload regulation, temporal organization, behavioral engagement and physiological stress, and may thus induce distinct adaptations.

In healthy mice, VWR and TR already elicit distinct molecular response in skeletal muscle, including in genes regulating protein turnover (Schmitt *et al*., 2020). Whether such differences persist during cancer, when tumor–host interactions profoundly alter host physiology, remains poorly understood. This question is further complicated by the fact that exercise itself may slow tumor growth (Pedersen *et al*., 2016). Differences in host outcomes between paradigms may therefore partly reflect differences in tumor burden, which must be accounted for to identify direct effects of exercise on host tissues.

Moreover, cachexia extends beyond skeletal muscle to adipose tissue, liver metabolism, systemic inflammation, energy homeostasis and behavior (Baracos *et al*., 2018), whereas exercise studies have focused predominantly on skeletal muscle mass and function. Whether these other compartments respond to exercise as skeletal muscle does remains largely unexplored, and addressing this question requires an integrative approach.

The UN-KC-6141 cell line derived from a pancreatic tumor arising in a KrasG12D;Pdx1-Cre (KC) mouse forms tumors in immunocompetent syngeneic C57BL/6 mice (Torres *et al*., 2013), thereby preserving the immune and inflammatory components of exercise responses. However, the systemic alterations induced by this model, and their responsiveness to exercise, remain poorly characterized.

Here, we compared VWR and TR in this model, assessing tumor progression, body composition, skeletal muscle, adipose tissue, hepatic metabolism, systemic inflammation and behavior, and adjusting for tumor burden to distinguish direct effects of exercise from effects secondary to tumor control. We hypothesized that VWR and TR would differentially modulate the systemic manifestations of pancreatic cancer cachexia and that exercise responsiveness would vary across physiological compartments.

## Material and methods

### Ethical approval statement

All experimental procedures involving animals were approved by the local Ethics Committee for Animal Experimentation (approval no. 23634) and were conducted in accordance with European Directive 2010/63/EU on the protection of animals used for scientific purposes. Experiments were performed at the M2S animal facility (approval no. A3504734) and the ARCHE animal facility of BIOSIT (UAR 3480 CNRS/US 18 Inserm, Université de Rennes; approval no. B3523840), Rennes, France. The work is reported in accordance with the ARRIVE guidelines.

### Cell culture

The pancreatic adenocarcinoma UN-KC-6141 cell line derived from a KrasG12D;Pdx1-Cre (KC) mouse was obtained from Prof. S. K. Batra (Torres *et al*., 2013) via a material transfer agreement (University of Nebraska Medical Center, 986099 Nebraska Medical Center, Omaha, NE 68198-6099, USA – March 2019). Cells were cultured in DMEM (high glucose) containing 10% heat inactivated fetal bovine serum (FBS), L-Glutamine (2 mM), non-essential amino acids (1 mM) and Penicillin/Streptomycin (1 mg/ml) under humidified conditions with 5% CO₂ at 37°C. Cells were split 1:6 every week and culture medium was renewed every two days.

### Animals and experimental design

A total of 100 eight-week-old male C57BL/6J mice were obtained from Janvier Labs (Le Genest-Saint-Isle, France). Mice were housed individually under standard laboratory conditions with a 12-h light/12-h dark cycle and ad libitum access to water and standard laboratory chow. Individual housing allowed longitudinal assessment of individual food intake and in CaVWR mice of voluntary wheel-running activity.

Twenty mice received a subcutaneous injection of phosphate-buffered saline (PBS) and constituted the Healthy group, whereas 80 mice received a subcutaneous injection of 1 × 10⁶ UN-KC-6141 cells suspended in 100 µl PBS into the right flank. On day 9 after tumor-cell injection, when tumors were reliably measurable by caliper, tumor-bearing mice were allocated to inactive cancer control (CaCTL), voluntary wheel running (CaVWR) or treadmill running (CaTR) groups. Allocation was performed to obtain comparable distributions of body weight and tumor size across the three tumor-bearing groups at the beginning of the exercise intervention. No random allocation procedure was used.

Group-specific exclusion criteria were applied. One Healthy mouse was excluded because of insufficient body-weight gain and one mouse allocated to CaCTL was excluded because no measurable tumor developed. Four mice allocated to CaVWR were excluded because they did not use the running wheel on a daily basis (fewer than 1000 wheel revolutions per day during the first week of the intervention). One CaTR mouse was excluded as an outlier for tumor-free body-weight change (more than 2.5 SD above the group mean). The final numbers included in the study were therefore Healthy, n = 19; CaCTL, n = 29; CaVWR, n = 21 and CaTR, n = 24.

Body weight, food intake and tumor dimensions were monitored longitudinally throughout the experimental period. Food intake was determined daily by weighing the food provided to each individually housed mouse and mean daily intake from day 9 to the end of the protocol was used for group comparisons. Tumor dimensions were measured using a digital caliper, and tumor volume was calculated as length × width × height × 0.5236 and expressed in mm³. A loss of >20% of body weight or tumor ulceration constituted humane endpoints requiring removal of the individual animal from the experimental protocol.

All animals completing the experimental protocol were euthanized on day 33 by an overdose of ketamine (200 mg kg⁻¹) and xylazine (10 mg kg⁻¹) administered via intraperitoneal injection. Tumors were excised and weighed immediately after collection. Final tumor-free body weight was calculated by subtracting the mass of the excised tumor from body weight measured immediately before killing. Tumor-free body-weight change was calculated relative to body weight measured on the day of tumor-cell injection (day 0).

Because of the nature of the exercise interventions, investigators conducting the training protocols were aware of group allocation. Blood was collected at the time of killing for subsequent biochemical analyses. Tumors, skeletal muscles, adipose tissues, liver, spleen and heart were excised and weighed as appropriate. Tissue samples were either snap-frozen in liquid nitrogen or fixed in 4% paraformaldehyde according to subsequent analyses. Samples were identified using individual animal identification numbers rather than experimental group labels, allowing downstream analyses to be performed without knowledge of group allocation.

The overall experimental design is illustrated in Figure 1.

**Figure 1.**
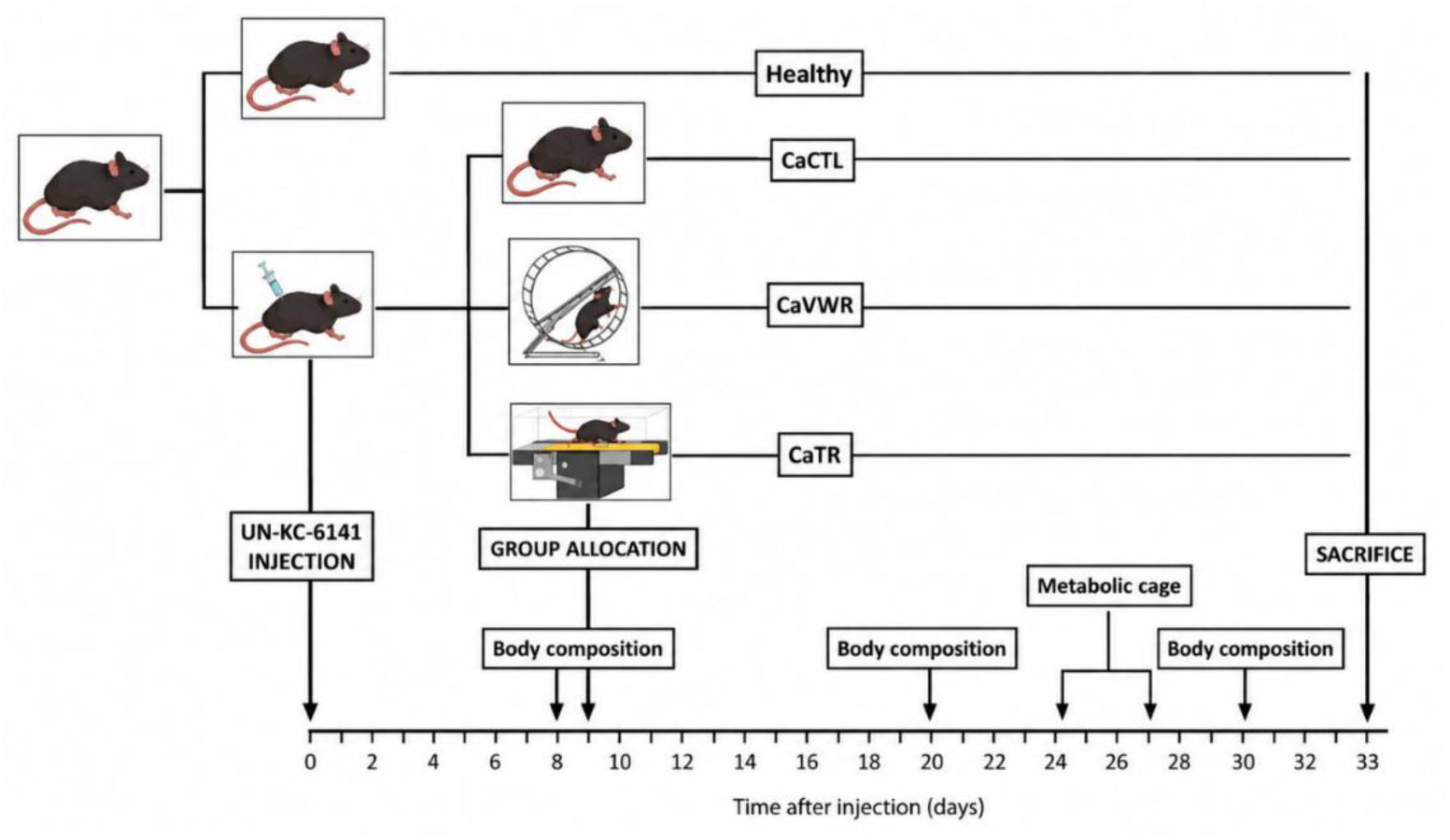
Study design.

### Exercise

Exercise interventions began on day 9 after tumor-cell injection and continued until the end of the experimental protocol. Two distinct exercise paradigms were investigated: voluntary wheel running (VWR) and treadmill running (TR).

### Voluntary wheel running

Mice allocated to the CaVWR group had continuous 24-h access to an individual running wheel in their home cage from day 9 onwards. The number of wheel revolutions was recorded each morning and converted into daily running distance according to wheel circumference. Running activity was entirely voluntary with no imposed duration, speed or timing of exercise. Mice that did not use the running wheel on a daily basis were excluded from the CaVWR group, as described above. Wheel access was discontinued 48 h before the end of the experimental protocol to limit the influence of acute exercise on terminal tissue and molecular measurements.

### Treadmill running

Mice allocated to the CaTR group underwent TR five days per week. Exercise sessions were performed during the late morning at a speed of 16 m min⁻¹ and a 10% incline. Exercise duration progressively increased from 20 min per session during the first week to 30 min per session during the second week and 40 min per session thereafter until the end of the intervention. Mice underwent treadmill familiarization before the training intervention. Animals were monitored throughout each session; predefined criteria specified that mice refusing to run or developing an exercise-related injury would be excluded from the CaTR group. No animals met these exclusion criteria.

### Forelimb muscle grip strength

Forelimb grip strength was assessed at the end of the experimental protocol using a 47200 Grip Strength Meter (Ugo Basile, Gemonio, Italy). Each mouse performed five consecutive trials. The three highest values were retained, and their mean was used for analysis (Dieumegard *et al*., 2025).

### Body composition

Body composition was assessed on days 8, 20 and 30 after tumor-cell injection using a Bruker Minispec system (Bruker, Germany). Lean mass, fat mass and body fluid were measured in conscious, non-fasted mice. Measurements were performed at fixed times of day to minimize variability related to circadian rhythms.

### Dietary behavior, exercise and energy expenditure

Whole-body energy metabolism, spontaneous locomotor activity, feeding behavior and sleep were assessed using a Promethion Core® metabolic phenotyping system from days 24 to 27 after tumor-cell injection. Five to six mice per group (Healthy and CaTR, n = 6; CaCTL and CaVWR, n = 5) were selected to obtain subsets representative of their respective groups in terms of body weight and, for tumor-bearing animals, tumor size. Mice were individually housed in the metabolic cages for four consecutive days at an ambient temperature of approximately 23.6 °C. The first two days were used for acclimatization to the metabolic cages. Data from the third day, corresponding to the first complete 24-h period after acclimatization and to a scheduled rest day for CaTR mice, were retained for analysis. All variables were analyzed separately during the light and dark phases.

Oxygen consumption (VO₂) and carbon dioxide production (VCO₂) were continuously measured and energy expenditure (EE) was calculated using the Weir equation. Resting energy expenditure was estimated as the mean EE during the 30-min period with the lowest activity level. Spontaneous locomotor activity and feeding behavior were continuously recorded by the Promethion system. Feeding variables included the number of feeding events and the amount of food consumed per feeding event. Sleep was estimated from patterns of inactivity continuously recorded by the Promethion system using its integrated sleep detection algorithm. During metabolic phenotyping, CaVWR mice retained continuous access to a running wheel, allowing voluntary wheel-running activity to be recorded separately from spontaneous cage locomotion. No running wheel was available to Healthy, CaCTL or CaTR mice.

### Cachexia severity classification

Cachexia severity was assessed at the end of the experimental protocol using a composite score based on tumor-free body weight, gastrocnemius muscle mass and epididymal adipose tissue mass. For each variable, the deviation from the Healthy group mean was expressed in standard deviation (SD) units. These standardized deviations were then averaged for each mouse to obtain an individual composite cachexia severity score. Mice were classified as having mild cachexia when the mean deviation was within 1 SD of the Healthy mean, moderate cachexia when the mean deviation was between 1 and 2 SD below the Healthy mean, and severe cachexia when the mean deviation was more than 2 SD below the Healthy mean, as previously described (Assi *et al*., 2016).

### Immunohistochemical analysis

*Gastrocnemius* muscle and epididymal adipose tissue samples were fixed in 4% paraformaldehyde, embedded in paraffin, and cut into 4-µm-thick sections.

Gastrocnemius sections were stained with CF640R-conjugated wheat germ agglutinin (WGA; Biotium, Fremont, CA, USA) to delineate muscle fiber boundaries. Muscle fiber cross-sectional area (CSA) was determined from 5 fields per tissue.

Epididymal adipose tissue sections were stained with hematoxylin-eosin-saffron (HES) and adipocyte cross-sectional area was determined from 5 fields per tissue. The sections were also immunostained for uncoupling protein 1 (UCP1) using an anti-UCP1 antibody (Abcam, Cambridge, UK; ab10983; 1:500). UCP1-positive staining area was quantified and expressed as a percentage of the total tissue area analyzed (UCP1-positive area/total tissue area × 100). All images were acquired using a Nanozoomer microscope (Hamamatsu, Shizuoka, Japan) at ×40 magnification. Muscle fiber and adipocyte CSA were quantified using ImageJ, whereas UCP1-positive area was quantified using QuPath.

### RNA extraction and RT-qPCR

RNA was extracted from tissue samples (*gastrocnemius*, liver) using Trizol reagent (Sigma Aldrich, St Louis, MO, USA). RNA concentration and quality were assessed, and reverse transcription was performed on 1 μg of total RNA. Real-time PCR was conducted with SybrGreen Supermix (Bio-Rad, Hercules, CA, USA) and specific primers, normalized to reference genes (Rpl4, Rpl19, Hprt), and analyzed using the ΔΔCt method in CFX-Manager software (Bio-Rad, Hercules, CA, USA).

**TABLE 1:**
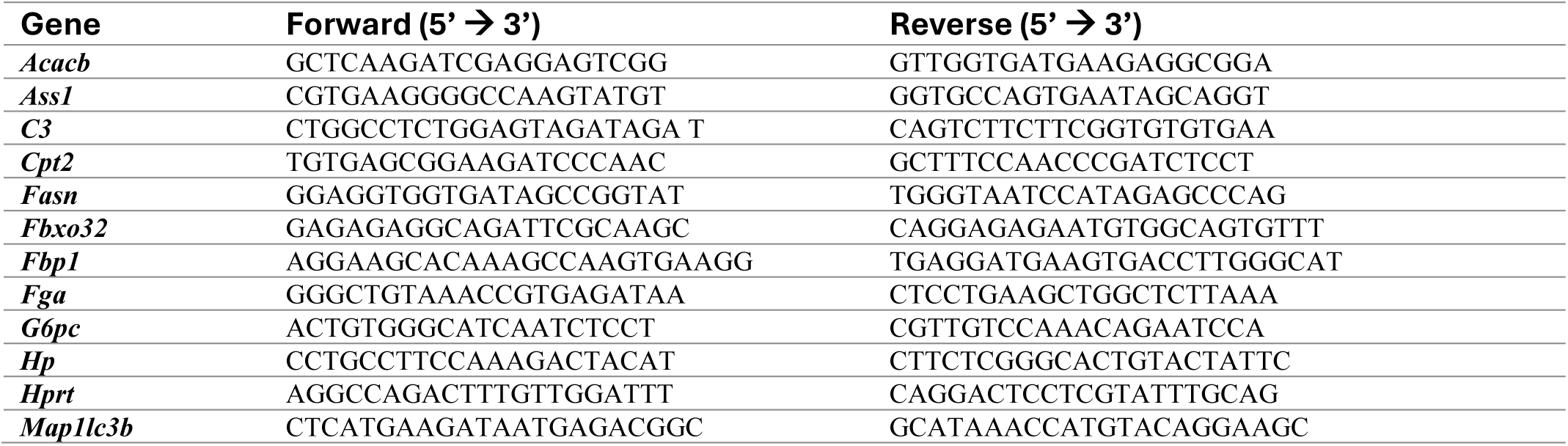
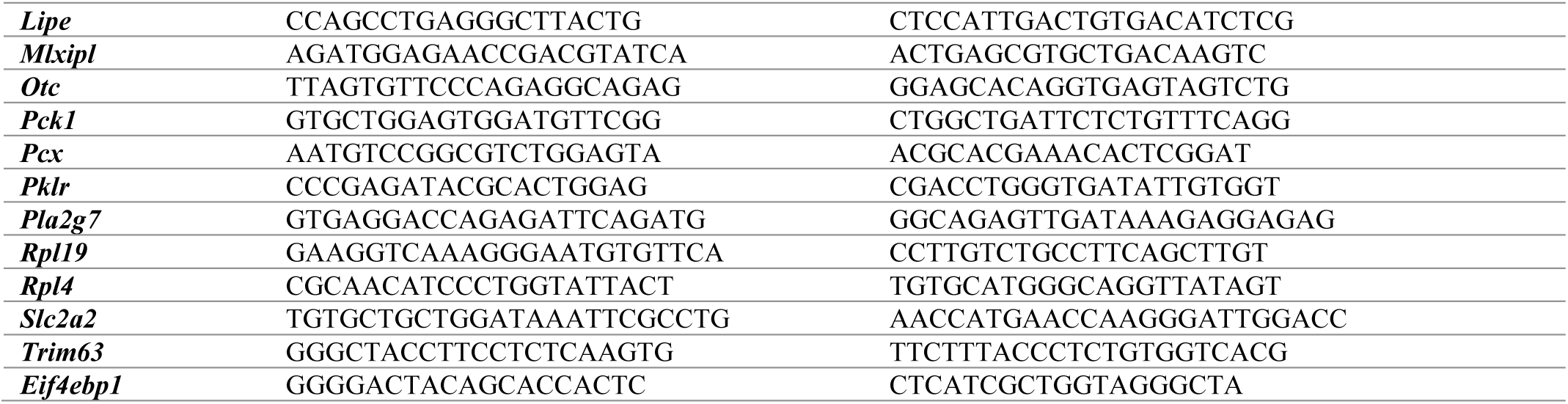
List of primers used for RT-qPCR analysis.

### Western blot analyses

*Gastrocnemius* muscle samples were homogenized in CHIP (50 mM Tris-HCl, 10 mM EDTA, 1% SDS, 50 mM NaF, 100 µM sodium orthovanadate) lysis buffer supplemented with protease and phosphatase inhibitors. Protein concentration was determined using the Lowry assay and equal amounts of protein (50 µg) were separated by SDS-PAGE and transferred onto nitrocellulose membranes (Bio-Rad, Hercules, CA, USA). Membranes were blocked with nonfat dry milk in TBS-Tween (0.05%), and incubated overnight at 4°C with primary antibodies against phosphorylated AKT (p-AKT; Cell Signaling Technology, catalog no. 9271; 1:1000), total AKT (Cell Signaling Technology, catalog no. 9272; 1:1000), and HSC70 (Santa Cruz; catalog no. sc-7298; 1:5000). After incubation with the appropriate secondary antibodies, proteins were detected using the Odyssey Infrared Imaging System (LI-COR Biosciences, Lincoln, NE, USA) and quantified by densitometry using ImageJ (U.S. National Institutes of Health, Bethesda, MD, USA) software. AKT phosphorylation was expressed as the p-AKT/total AKT ratio. HSC70 was used as a loading control.

### Enzyme-linked immunosorbent assay

Plasma IL-6 concentrations were measured using a commercially available mouse IL-6 ELISA kit (LEGEND MAX™ Mouse IL-6 ELISA Kit, BioLegend) according to the manufacturer’s instructions. Samples were analyzed in duplicate, and absorbance was measured at 450 nm using a SpectraMax Plus 384 microplate reader (Molecular Devices, San Jose, CA, USA). IL-6 concentrations were calculated from the corresponding standard curve and expressed as pg ml⁻¹.

### Hepatic phospholipid analysis

Lipids were extracted as previously described (Bligh & Dyer, 1959). In order to quantify liver mitochondrial phospholipids (cardiolipin [CL], phosphatidylethanolamine [PE], phosphatidylinositol [PI], phosphatidylcholine [PC], phosphatidylserine [PS] and sphingomyelin [SM]), standards and samples were loaded on silica plates using Linomat V sample applicator (CAMAG, Muttenz Switzerland). After migration, the plates were immersed into 10% copper sulfate (wt/vol) in 8% phosphoric acid solution (vol/vol). Plates were heated (160 °C for 15 min) to stain phospholipids. Quantification was performed using TLC-visualizer Reprostar 3® and WinCats VideoScan® software (CAMAG, Muttenz, Switzerland). Results were expressed in µg of phospholipids per µg of protein.

### Statistical analyses

Data are presented as mean ± SD. Statistical analyses were performed using GraphPad Prism version 8.0.1 (GraphPad Software, San Diego, CA, USA), except for ANCOVA analyses, which were performed using R (version 4.6.1).

Normality of data distribution was assessed using the Shapiro-Wilk test. For comparisons between experimental groups at a single time point, normally distributed data were analyzed using one-way ANOVA followed by Dunnett’s multiple-comparison test, whereas non-normally distributed data were analyzed using the Kruskal-Wallis test followed by Dunn’s multiple-comparison test.

Longitudinal data, including body weight, tumor growth, food intake and body-composition measurements, were analyzed using linear mixed-effects models with group, time and group × time interaction as fixed effects and individual mouse as a random effect.

To determine whether differences in tissue mass between exercise groups could be explained solely by differences in tumor burden, analyses of covariance (ANCOVA) were performed using tissue mass as the dependent variable, experimental group as the fixed factor and final tumor mass as the covariate. The group x tumor mass interaction was initially tested to assess the homogeneity of regression slopes. When the interaction was not significant, the model was refitted without the interaction term and adjusted group effects were assessed. Estimated marginal means were subsequently compared between groups with Tukey adjustment for multiple comparisons. ANCOVA analyses were performed separately for gastrocnemius muscle mass and epididymal adipose tissue mass. Within the CaVWR group, associations between mean daily running distance and gastrocnemius mass or final tumor-free body weight were assessed using Pearson’s correlation. Within tumor-bearing mice that underwent metabolic phenotyping, associations between estimated dark-phase sleep duration and tumor-free body-weight change or epididymal adipose tissue mass were assessed using Spearman’s rank correlation and multiple linear regression including experimental group and tumor mass as covariates.

Because several analyses were performed on subsets of animals or tissue samples, sample sizes vary between analyses and are reported for each panel in the figure legends. Comparisons were considered significant for P values below 0.05. Exact P values are reported whenever possible.

## RESULTS

### UN-KC-6141 pancreatic tumors induce a systemic cachectic phenotype that is only partially modified by voluntary wheel running

To establish the systemic phenotype induced by UN-KC-6141 pancreatic tumors, tumor-bearing mice (CaCTL) were first compared with Healthy mice. Tumor volume increased progressively in all tumor-bearing groups throughout disease progression (Fig. 2A). CaCTL mice exhibited significantly reduced tumor-free body-weight gain (0.34 ± 6.02% vs. 8.13 ± 4.35%) compared with the Healthy group (Fig. 2B), fat mass loss (−6.43 ± 10.25% vs. +6.69 ± 7.59%) (Fig. 2C), a non-significant reduction in lean mass accretion, although this measurement includes the tumor (3.47 ± 3.95% vs. 5.26 ± 2.38%) (Fig. 2C), together with splenomegaly and elevated circulating IL-6 concentrations (Fig. 2E–F). These alterations confirmed the development of a systemic cachectic phenotype in UN-KC-6141 tumor-bearing mice.

**Figure 2.**
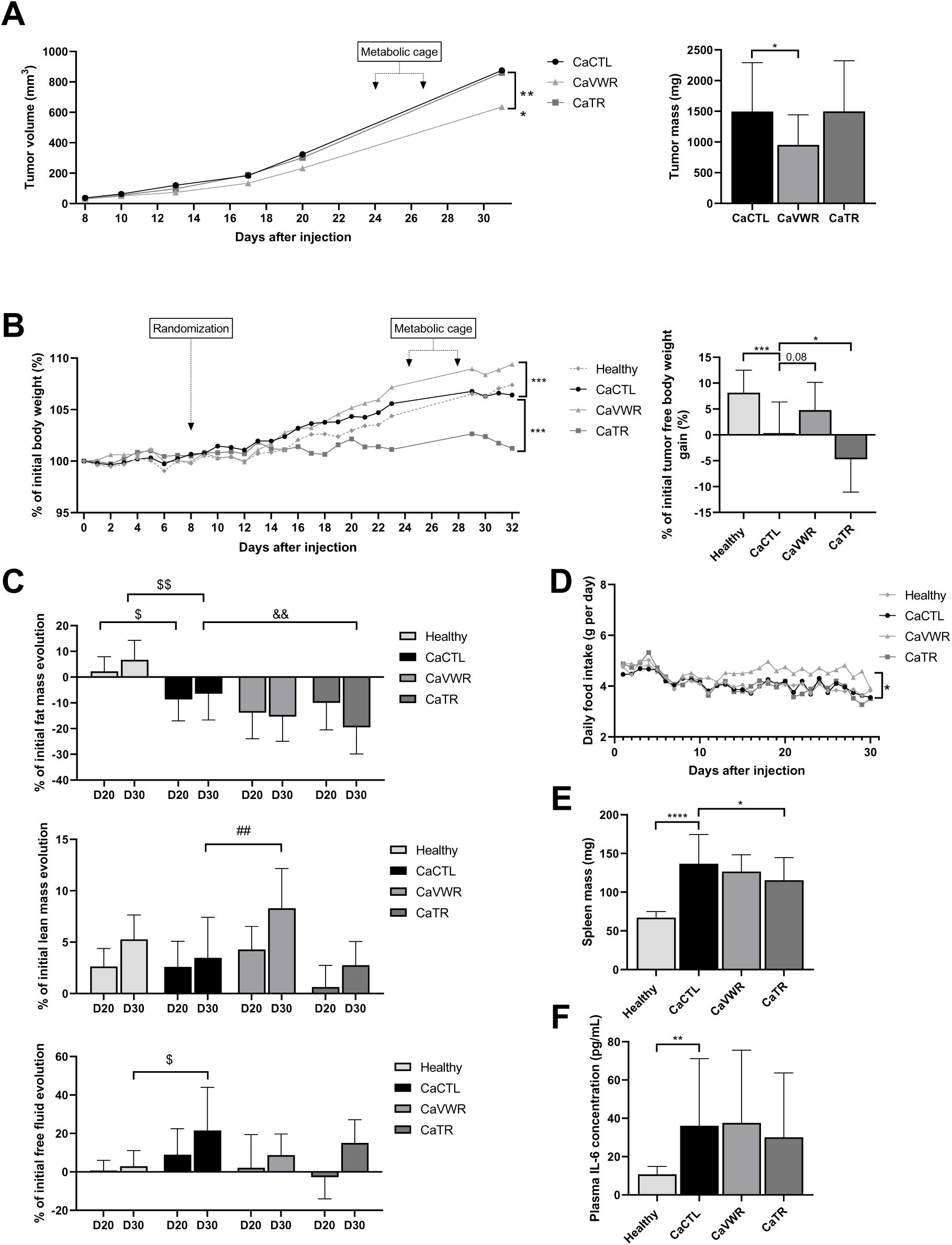
UN-KC-6141 pancreatic tumors induce a systemic cachectic phenotype that is only partially modified by voluntary wheel running. Male C57BL/6J mice were allocated to Healthy, inactive tumor-bearing (CaCTL), voluntary wheel running (CaVWR) or treadmill running (CaTR) groups; exercise interventions started on day 9 after tumor-cell injection. VWR slowed tumor growth, reduced final tumor mass and preserved lean mass, whereas neither exercise paradigm prevented fat mass loss, fluid accumulation or the systemic inflammatory phenotype. (A) Longitudinal tumor growth and final tumor mass. (B) Longitudinal changes in body weight and final tumor-free body-weight change. (C) Longitudinal changes in body composition, including fat mass, lean mass and free fluid (low-field NMR; the lean and fluid compartments include the tumor). (D) Longitudinal daily food intake (E) Spleen mass at the end of the experimental protocol. (F) Plasma IL-6 concentration. Data are presented as mean ± SD. Healthy, n = 19; CaCTL, n = 29; CaVWR, n = 21; CaTR, n = 24, except for body composition (C; n = 10, 14, 7 and 10, respectively) and plasma IL-6 (F; n = 9, 14, 14 and 15, respectively). Longitudinal variables were analyzed using linear mixed-effects models, whereas comparisons at a single time point were performed using one-way ANOVA followed by Dunnett’s multiple-comparison test or Kruskal–Wallis followed by Dunn’s multiple-comparison test, as appropriate. In (A), (B) and (D), asterisks next to the curves indicate a significant group × time interaction versus CaCTL (linear mixed-effects model; \**P* < 0.05, \*\**P* < 0.01, \*\*\**P* < 0.001). In bar graphs, brackets indicate significant differences (\**P* < 0.05, \*\**P* < 0.01, \*\*\**P* < 0.001, \*\*\*\**P* < 0.0001); exact P values are shown for non-significant differences with 0.05 ≤ *P* < 0.10. In (C), $*P* < 0.05 and $$*P* < 0.01, Healthy vs. CaCTL; #*P* < 0.05 and ##*P* < 0.01, CaVWR vs. CaCTL; &&P < 0.01, CaTR vs. CaCTL, at the same time point.

We next investigated whether this systemic phenotype was similarly modified by two commonly used aerobic exercise modalities. While tumor volume increased progressively in all tumor-bearing groups (Fig. 2A), VWR slowed tumor growth during the late stages of disease progression compared with inactive tumor-bearing mice (group × time interaction, *P* < 0.001) and reduced tumor mass at dissection (CaVWR: 951.9 ± 490.4 vs. CaCTL: 1494.9 ± 797.2 mg, P = 0.025). In contrast, TR did not modify tumor growth kinetics or final tumor burden.

These differences in tumor progression were accompanied by divergent body-weight trajectories. Longitudinal analysis showed that VWR significantly improved body-weight evolution compared with CaCTL, whereas TR significantly worsened it (Fig. 2B). At the final time point, CaVWR mice showed greater tumor-free body-weight gain than CaCTL mice (+4.78 ± 5.36% vs. +0.34 ± 6.02%), although this difference did not reach statistical significance (*P* = 0.081), whereas CaTR mice showed a significant reduction in tumor-free body weight (−4.71 ± 6.36%, P = 0.037) compared with CaCTL mice. Daily food intake did not differ between Healthy and CaCTL mice (4.01 ± 0.33 vs. 4.04 ± 0.43 g day⁻¹; P > 0.99), indicating that the cachectic phenotype developed without a reduction in food intake. In contrast, from the onset of the intervention, CaVWR mice (4.57 ± 0.65 g day⁻¹; P = 0.020) ate approximately 13% more than CaCTL mice whereas food intake was unchanged in CaTR mice (4.04 ± 0.59 g day⁻¹; P > 0.99; Fig. 2D).

Longitudinal body composition analysis revealed distinct temporal patterns across physiological compartments (Fig. 2C). In tumor-bearing mice, fat mass loss was already significant at day 20 (P = 0.020), whereas lean mass was only marginally affected, possibly because the low-field NMR lean compartment includes the growing tumor, indicating that the different components of the cachectic phenotype did not develop simultaneously. We next examined whether these compartment-specific trajectories were similarly affected by exercise. VWR significantly preserved lean mass (+8.30 ± 3.87%) compared with inactive tumor-bearing mice (+3.47 ± 3.95%, P = 0.008), whereas TR exercise had no detectable effect (+2.75 ± 2.30%). In contrast, neither exercise modality prevented fat mass wasting which reached −15.35 ± 9.57% in CaVWR and −19.46 ± 10.41% in CaTR mice, fat mass loss being even greater in CaTR than in CaCTL mice at the final time point (P = 0.007). Likewise, fluid accumulation progressively increased during tumor progression (P = 0.020 at the final time point) and remained unaffected by either exercise intervention.

To determine whether exercise also modified the systemic inflammatory response associated with tumor progression, spleen mass and circulating IL-6 concentrations were measured at dissection (Fig. 2E–F). However, VWR did not significantly reduce spleen mass or plasma IL-6 concentrations, and TR only modestly reduced spleen mass (115.4 ± 29.3 vs. 136.6 ± 38.1 mg, *P* = 0.022) without affecting IL-6, indicating persistent systemic inflammation despite exercise.

Collectively, these findings show that VWR modified some components of the cachectic phenotype with slower tumor growth and lower final tumor mass, a more favorable body-weight trajectory and greater lean mass accretion, whereas the TR protocol used in this study did not reproduce these adaptations and even significantly exacerbated body-weight loss.

### Voluntary wheel running selectively preserves skeletal muscle after adjustment for tumor burden

We next investigated whether skeletal muscle represented a preferentially protected tissue during pancreatic tumor progression. Compared with Healthy mice, tumor-bearing animals (CaCTL) exhibited marked muscle atrophy, as demonstrated by reduced gastrocnemius and heart masses (the reduction in soleus mass did not reach statistical significance; *P* = 0.063), together with a significant decrease in gastrocnemius fiber cross-sectional area (Fig. 3A–C). VWR significantly attenuated these alterations, increasing gastrocnemius mass (CaVWR: 149.58 ± 17.60 vs. CaCTL: 128.01 ± 21.54 mg, P = 0.001), soleus mass (CaVWR: 8.21 ± 0.99 vs. CaCTL: 6.74 ± 1.25 mg, *P* < 0.0001) compared with CaCTL mice, whereas heart mass was not significantly modified (CaVWR: 121.6 ± 13.08 vs. CaCTL: 114.45 ± 18.33 mg, *P* = 0.131). Gastrocnemius fiber cross-sectional area did not differ significantly between CaVWR and CaCTL mice (940.25 ± 119.16 vs. 816.28 ± 175.29 µm², *P* = 0.213). In contrast, TR exercise failed to improve any morphological indices of skeletal muscle preservation, all remaining comparable to those observed in inactive tumor-bearing mice.

**Figure 3.**
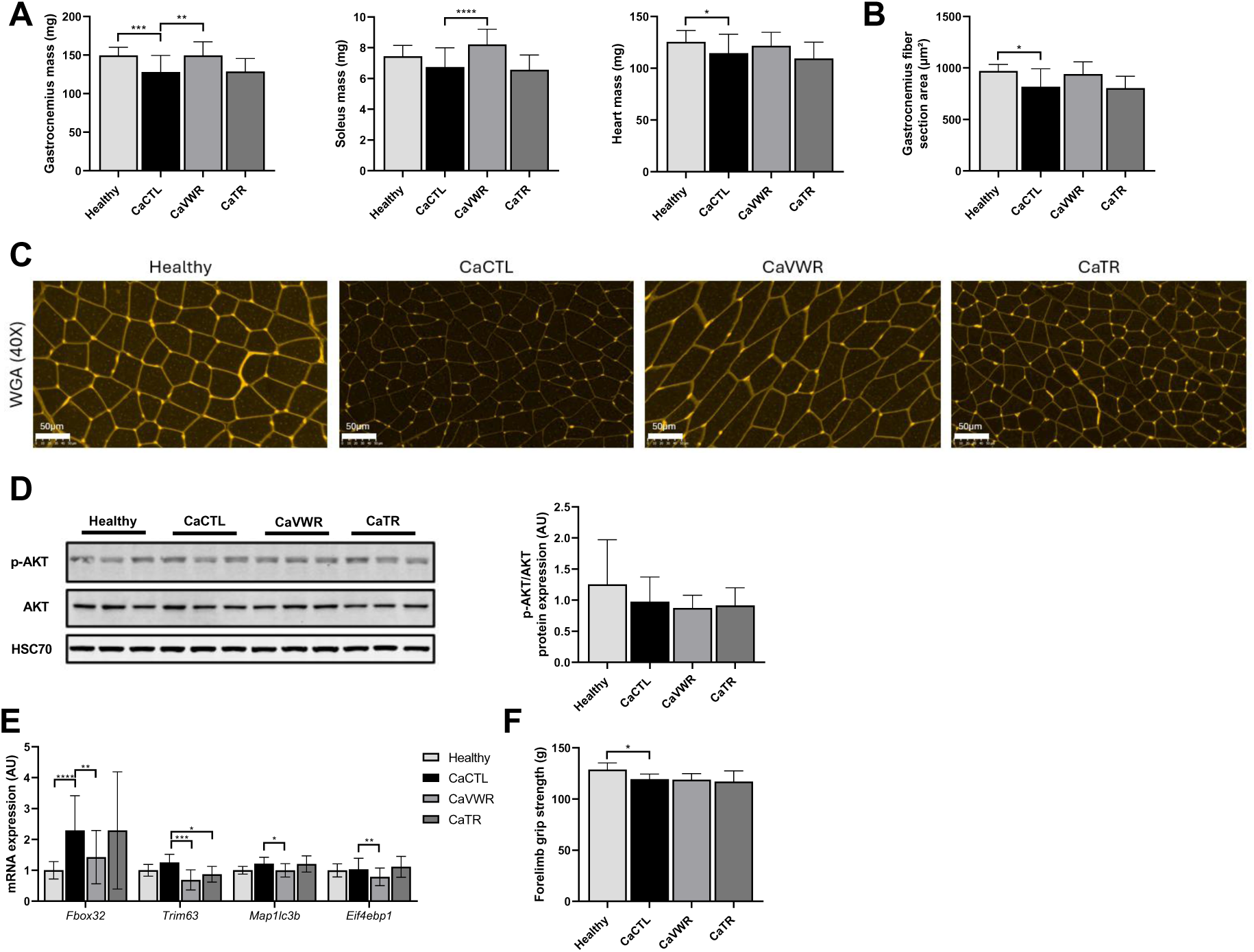
Voluntary wheel running selectively preserves skeletal muscle during pancreatic cancer cachexia. VWR preserved skeletal muscle mass and was associated with lower expression of catabolic and autophagy-related markers, whereas muscle preservation was not accompanied by restoration of grip strength. (A) Gastrocnemius, soleus and heart masses at the end of the experimental protocol. (B) Gastrocnemius muscle fiber cross-sectional area. (C) Representative gastrocnemius muscle sections stained with wheat germ agglutinin (WGA). (D) Representative western blots of p-AKT, AKT and HSC70 (three animals per group) and AKT phosphorylation expressed as the p-AKT/total AKT ratio. (E) Gastrocnemius expression of the muscle protein-turnover-related genes *Fbxo32*, *Trim63*, *Map1lc3b* and *Eif4ebp1*. (F) Forelimb grip strength. Data are presented as mean ± SD. Muscle and heart masses (A): Healthy, n = 19; CaCTL, n = 29; CaVWR, n = 21; CaTR, n = 24. Fiber cross-sectional area (B): n = 10, 14, 7 and 7; AKT phosphorylation (D): n = 9, 16, 14 and 14; gene expression (E): n = 9–19, 12–25, 12–19 and 13–23; grip strength (F): n = 9, 15, 14 and 14, for Healthy, CaCTL, CaVWR and CaTR, respectively. Comparisons were performed using one-way ANOVA followed by Dunnett’s multiple-comparison test or Kruskal–Wallis followed by Dunn’s multiple-comparison test, as appropriate. Brackets indicate significant differences (\**P* < 0.05, \*\**P* < 0.01, \*\*\**P* < 0.001, \*\*\*\**P* < 0.0001); exact P values are shown for non-significant differences with 0.05 ≤ *P* < 0.10.

Because VWR also reduced tumor burden (Fig. 2), we next determined whether the greater skeletal muscle mass observed in CaVWR mice could be explained solely by differences in tumor burden. An analysis of covariance (ANCOVA) using tumor mass as a covariate demonstrated a strong association between tumor mass and gastrocnemius mass (tumor mass effect: *P* < 0.0001). Importantly, a significant effect of experimental group remained after adjustment for tumor mass (group effect: *P* = 0.013), whereas the tumor mass × group interaction was not significant (*P* = 0.633) indicating similar regression slopes across groups. Estimated marginal means further revealed that gastrocnemius mass remained significantly greater in CaVWR mice than in both CaCTL (*P* = 0.016) and CaTR (*P* = 0.030) mice after adjustment for tumor burden. Thus, although gastrocnemius mass was strongly associated with tumor burden, the effect of VWR on skeletal muscle mass persisted after adjustment for tumor burden.

Running distance varied substantially within the group of CaVWR mice (4.03 ± 2.32 km day⁻¹; range 0.32–7.62 km day⁻¹). However, mean daily running distance was not associated with gastrocnemius mass (r = −0.15, P = 0.51; Fig. S1A) or final tumor-free body weight (r = −0.25, P = 0.28; Fig. S1B).

To determine whether these structural adaptations were associated with altered muscle protein turnover, we next quantified key molecular regulators of anabolic and catabolic signaling (Fig. 3D–E). Tumor progression significantly increased the expression of the atrophy-related gene Fbxo32 in gastrocnemius, whereas Trim63 and Map1lc3b expression was not significantly altered (P = 0.434 and P = 0.072, respectively) and AKT phosphorylation remained unchanged. VWR significantly reduced *Fbxo32*, *Trim63*, *Eif4ebp1* and *Map1lc3b* expression compared with CaCTL mice. Apart from a lower Trim63 expression (P = 0.027), these changes were not observed in CaTR mice.

Despite these structural and molecular adaptations, forelimb grip strength remained significantly reduced in tumor-bearing mice compared with healthy mice (CaCTL: 119.44 ± 5.02 vs. Healthy: 128.86 ± 6.48 g, P = 0.010) and was not restored by either exercise (Fig. 3F). Together, these findings identify skeletal muscle as one of the physiological compartments most responsive to VWR exercise during pancreatic cancer cachexia.

### Adipose tissue and hepatic metabolic remodeling remain largely resistant to exercise during pancreatic cancer cachexia

Having established that skeletal muscle was responsive to VWR, we next determined whether similar adaptations occurred in other major metabolic tissues involved in the cachectic syndrome. Consistent with the progressive alterations in body composition observed longitudinally (Fig. 2C), pancreatic tumor progression induced a marked reduction in epididymal adipose tissue mass (CaCTL: 199.85 ± 108.14 vs. Healthy: 292.7 ± 88.5 mg, P = 0.002) together with a pronounced decrease in adipocyte cross-sectional area (CaCTL: 2529.32 ± 907.48 vs. Healthy: 3996.27 ± 879.67 µm², P = 0.001), confirming substantial adipose tissue wasting in tumor-bearing mice (Fig. 4A–B). In contrast to skeletal muscle, neither VWR nor TR attenuated these alterations, as epididymal adipose tissue mass and adipocyte morphology remained comparable to those observed in CaCTL tumor-bearing animals (Fig. 4A–B). Because VWR reduced tumor burden, we next examined whether the absence of adipose tissue preservation could simply reflect differences in tumor size. ANCOVA using tumor mass as a covariate revealed a strong negative association between tumor burden and adipose tissue mass (*P* < 0.0001). However, unlike skeletal muscle, adjustment for tumor mass did not reveal a protective effect of voluntary wheel running. Rather, at a given tumor mass, adipose tissue mass was significantly lower in CaVWR than in CaCTL mice (Tukey-adjusted *P* = 0.005), and a similar reduction was observed in CaTR mice (*P* = 0.040), indicating that exercise was associated with greater adipose tissue loss at a given tumor burden.

**Figure 4.**
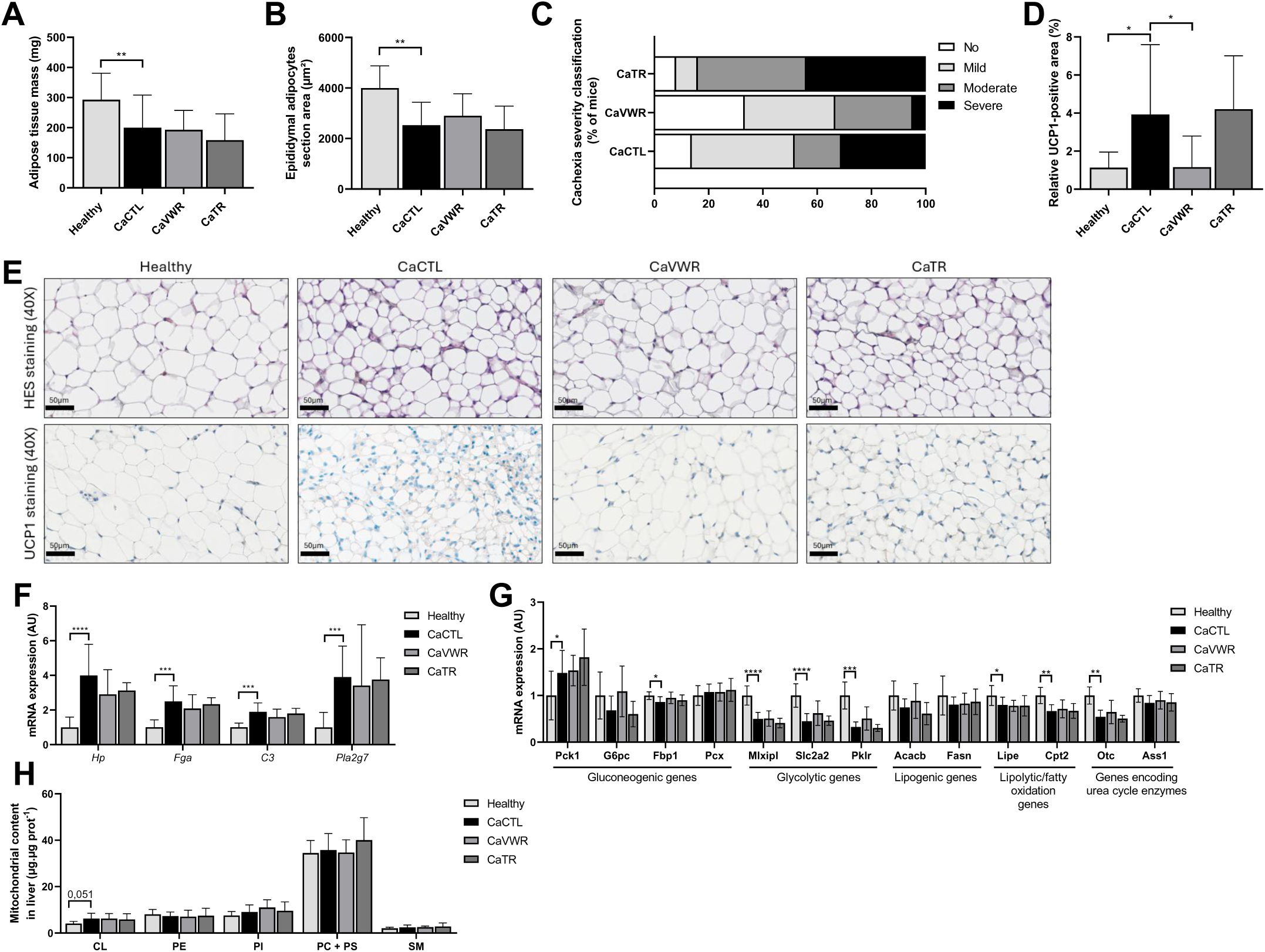
Adipose tissue and hepatic metabolic remodeling remain largely resistant to exercise during pancreatic cancer cachexia. Pancreatic tumor progression induced marked adipose tissue wasting and hepatic metabolic remodeling. In contrast to skeletal muscle, these alterations were largely unaffected by either exercise paradigm. (A) Epididymal adipose tissue mass at the end of the experimental protocol. (B) Adipocyte cross-sectional area (HES staining; representative sections in E). (C) Composite cachexia severity classification based on tumor-free body weight, gastrocnemius muscle mass and epididymal adipose tissue mass, expressed as the percentage of mice in each category; mice were classified according to their mean standardized deviation from Healthy mice. (D) Relative UCP1-positive area. (E) Representative adipose tissue sections stained with HES (top) and immunostained for UCP1 (bottom). (F) Hepatic expression of acute-phase and inflammatory response genes (*Hp*, *Fga*, *C3* and *Pla2g7*). (G) Hepatic expression of genes involved in glucose, lipid and amino-acid metabolism. (H) Hepatic phospholipid content, including cardiolipin (CL). Data are presented as mean ± SD. Adipose tissue mass (A): Healthy, n = 19; CaCTL, n = 29; CaVWR, n = 21; CaTR, n = 24. Adipocyte cross-sectional area (B): n = 10, 14, 7 and 10; UCP1 (D): n = 13, 14, 7 and 11; hepatic gene expression (F, G): n = 9–10, 12–13, 8–9 and 10 (*Fasn*, Healthy n = 4); hepatic phospholipids (H): n = 8–9, 13–14, 13 and 14, for Healthy, CaCTL, CaVWR and CaTR, respectively. Comparisons were performed using one-way ANOVA followed by Dunnett’s multiple-comparison test or Kruskal–Wallis followed by Dunn’s multiple-comparison test, as appropriate. Brackets indicate significant differences (\**P* < 0.05, \*\**P* < 0.01, \*\*\**P* < 0.001, \*\*\*\**P* < 0.0001); exact P values are shown for non-significant differences with 0.05 ≤ *P* < 0.10.

Finally, we examined the overall severity of the cachectic phenotype by integrating tumor-free body weight, gastrocnemius muscle mass and epididymal adipose tissue mass into a composite cachexia severity classification (Fig. 4C). CaVWR mice descriptively showed a shift toward milder cachexia compared with CaCTL mice. Importantly, this composite response did not reflect uniform protection across tissues, as skeletal muscle preservation occurred despite persistent adipose tissue wasting. Interestingly, the relative UCP1-positive area was greater in CaCTL than in Healthy mice (CaCTL: 3.92 ± 3.66% vs. Healthy: 1.12 ± 0.82%, P = 0.034) and was significantly lower in CaVWR mice (1.15 ± 1.63%, P = 0.034 vs. CaCTL), whereas no difference was observed between CaTR (4.20 ± 2.80%) and CaCTL mice (Fig. 4D–E).

We next investigated whether the limited responsiveness to exercise observed in adipose tissue also extended to the liver. Compared with Healthy mice, CaCTL mice exhibited marked hepatic transcriptional remodeling, characterized by increased expression of acute-phase response genes, including *Hp* (3.99 ± 1.80 AU), *Fga* (2.50 ± 0.90 AU), *C3* (1.89 ± 0.52 AU) and *Pla2g7* (3.91 ± 1.78 AU), together with alterations in genes involved in glucose and lipid metabolism (Fig. 4F–G). These included increased expression of Pck1 and decreased expression of Mlxipl, Slc2a2, Pklr, Fbp1, Lipe, Cpt2 and Otc. Neither VWR nor TR substantially restored these hepatic transcriptional alterations, with expression profiles remaining largely comparable to those observed in CaCTL mice.

Hepatic mitochondrial phospholipid composition was also examined (Fig. 4H). Cardiolipin content was 53% higher in liver of tumor-bearing mice, although this difference did not reach statistical significance (CaCTL: 6.24 ± 2.28 vs. Healthy: 4.09 ± 0.91 µg µg^-1^ protein, *P* = 0.051) and was not significantly modified by either VWR or TR.

Together, these findings indicate that, unlike skeletal muscle, adipose tissue was not preserved by either exercise paradigm, despite changes in adipose tissue responses, while hepatic metabolic alterations remained largely unchanged.

### Voluntary wheel running and treadmill running differentially alter behavioral and metabolic organization during pancreatic cancer cachexia

Having demonstrated that voluntary wheel running selectively preserves skeletal muscle without restoring systemic metabolic alterations, we next investigated whether the two exercise paradigms were associated with distinct behavioral and metabolic responses during pancreatic cancer cachexia (Fig. 5).

**Figure 5.**
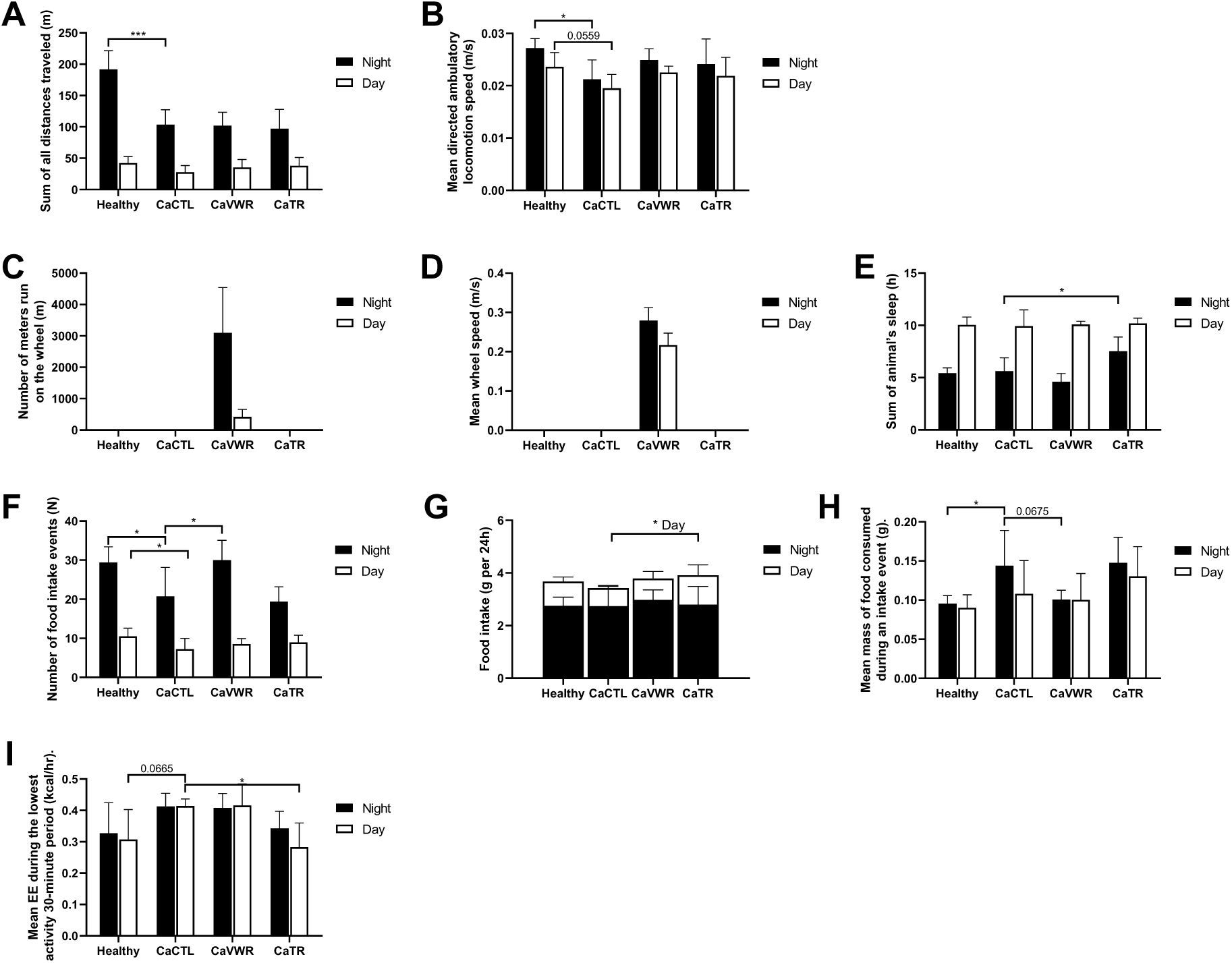
Voluntary wheel running and treadmill running are associated with distinct behavioral and metabolic patterns during pancreatic cancer cachexia. Whole-body activity, voluntary wheel-running activity, feeding behavior, sleep and resting energy expenditure were assessed using metabolic phenotyping. VWR increased voluntary activity without a detectable compensatory reduction in spontaneous cage locomotion and was associated with changes in feeding organization, whereas TR was associated with increased dark-phase sleep duration and reduced light-phase resting energy expenditure. (A) Spontaneous distance traveled during the dark (night) and light (day) phases. (B) Mean spontaneous locomotion speed. (C) Voluntary wheel-running distance in CaVWR mice. (D) Mean wheel-running speed in CaVWR mice. (E) Estimated sleep duration. (F) Number of feeding events. (G) Food intake during the dark (night, black) and light (day, white) phases; stacked bars show total 24-h intake, and error bars indicate the SD of each phase. (H) Amount of food consumed per feeding event. (I) Resting energy expenditure. Data are presented as mean ± SD. Metabolic phenotyping was performed in five to six mice per group (Healthy and CaTR, n = 6; CaCTL and CaVWR, n = 5) selected as representative of their respective experimental groups. Comparisons were performed separately for the dark and light phases using one-way ANOVA followed by Dunnett’s multiple-comparison test versus CaCTL. Brackets indicate significant differences (*P < 0.05, ***P < 0.001); in (G), *P < 0.05, CaTR vs. CaCTL during the light phase, whereas total 24-h intake did not differ between groups (all P > 0.5); exact P values are shown for non-significant differences with 0.05 ≤ *P* < 0.10.

Tumor progression markedly altered spontaneous locomotor behavior. Compared with Healthy mice, CaCTL mice exhibited a marked reduction in total distance traveled during the dark phase (Healthy: 191.8 ± 29.7 m vs. CaCTL: 103.4 ± 24.0 m; *P* < 0.001), together with a lower locomotion speed during the dark phase (Healthy: 0.027 ± 0.002 vs. CaCTL: 0.021 ± 0.004 m s⁻¹; *P* = 0.024), whereas the difference during the light phase did not reach statistical significance (*P* = 0.0569). Neither VWR nor TR significantly modified these parameters compared with CaCTL mice during either phase (all *P* > 0.2; Fig. 5A–B).

Wheel-running activity was confirmed in CaVWR mice, which covered a mean distance of 3100.0 ± 1439.4 m during the dark phase and 418.9 ± 239.1 m during the light phase, at mean speeds of 0.279 ± 0.033 m s⁻¹ and 0.216 ± 0.031 m s⁻¹, respectively (Fig. 5C–D). Notably, spontaneous cage locomotion outside the wheel did not differ between CaVWR and CaCTL mice (Fig. 5A), indicating that voluntary wheel engagement was not accompanied by a compensatory reduction in spontaneous cage activity. When wheel-running and in-cage locomotion were considered together, CaVWR mice covered a greater total distance than CaCTL mice during both the dark phase (3201.9 vs. 103.4 m) and the light phase (454.2 vs. 27.7 m). At this stage of the intervention, CaTR mice were prescribed 40-min treadmill sessions at 16 m min⁻¹ and a 10% incline, corresponding to a theoretical running distance of approximately 640 m per session.

In contrast to locomotor activity, sleep organization differed across groups. CaTR mice exhibited a significant increase in estimated sleep duration during the dark phase compared with CaCTL mice (7.52 ± 1.36 h vs. 5.62 ± 1.28 h; *P* = 0.019), whereas CaVWR mice displayed a sleep pattern comparable to CaCTL mice (4.61 ± 0.79 h; *P* = 0.312). No differences were observed during the light phase for any group (all P > 0.9; Fig. 5E). Across tumor-bearing mice, dark-phase sleep duration was inversely correlated with tumor-free body-weight change (ρ = - 0.85, P < 0.001) and epididymal adipose tissue mass (ρ = −0.61, P = 0.013), and both associations remained significant after adjustment for group and tumor mass (P = 0.017 and P = 0.045, respectively). Feeding behavior was also differentially affected. CaCTL mice displayed fewer feeding events than Healthy mice during both the dark phase (20.7 ± 7.5 vs. 29.4 ± 4.0; *P* = 0.031) and the light phase (7.2 ± 2.8 vs. 10.5 ± 2.1; *P* = 0.043), together with a greater amount of food consumed per feeding event during the dark phase (0.144 ± 0.045 vs. 0.095 ± 0.010 g; *P* = 0.030), consistent with altered feeding organization. CaVWR mice displayed significantly more feeding events than CaCTL mice during the dark phase (30.0 ± 5.1 vs. 20.7 ± 7.5; *P* = 0.027), whereas the reduction in the amount of food consumed per feeding event did not reach statistical significance (0.101 ± 0.012 g; *P* = 0.068). In contrast, the number of feeding events and the amount of food consumed per feeding event did not differ between CaTR and CaCTL mice (all P > 0.37), whereas CaTR mice consumed more food than CaCTL mice during the light phase (1.12 ± 0.40 vs. 0.69 ± 0.07 g; P = 0.037; Fig. 5F–H). Total 24-h food intake, however, did not differ between groups (all P > 0.5), indicating a temporal redistribution of food intake toward the light phase in CaTR mice rather than an increase in total intake.

Resting energy expenditure was higher in CaCTL than in Healthy mice during the light phase, although this difference did not reach statistical significance (0.414 ± 0.022 vs. 0.307 ± 0.095 kcal h⁻¹; *P* = 0.066). Resting energy expenditure was significantly lower in CaTR than in CaCTL mice during the light phase (0.283 ± 0.077 vs. 0.414 ± 0.022 kcal h⁻¹; *P* = 0.022), whereas no significant difference was observed during the dark phase (0.343 ± 0.054 vs. 0.412 ± 0.042 kcal h⁻¹; *P* = 0.223). CaVWR mice remained comparable to CaCTL mice during both phases (both P > 0.99; Fig. 5I). Body mass recorded in the metabolic cages was only slightly lower in CaTR than in CaCTL mice (23.2 ± 0.8 vs. 23.9 ± 1.4 g, P = 0.76), consistent with their lower body-weight trajectory (Fig. 2B); this difference of approximately 3% cannot account for the 17–32% lower resting energy expenditure observed in CaTR mice. Finally, the increased duration of the dark-phase sleep, light-phase food intake and the reduced resting energy expenditure of CaTR mice were directionally consistent across the four complete 24-h periods recorded (data not shown), including the two days on which CaTR mice performed a training session.

Collectively, these findings indicate that VWR and TR were associated with distinct behavioral and metabolic patterns during pancreatic cancer cachexia. VWR was associated with changes in feeding organization, whereas TR was associated with longer dark-phase sleep duration, greater light-phase food intake and reduced light-phase resting energy expenditure.

## Discussion

Using an integrative physiological approach, the present study shows that exercise responsiveness during pancreatic cancer cachexia is tissue-specific: VWR preserved skeletal muscle despite persistent alterations in adipose tissue, hepatic metabolism and systemic inflammation. The greater muscle mass observed in CaVWR mice remained evident after adjustment for tumor burden, indicating that reduced tumor growth alone could not fully explain the preservation of skeletal muscle mass. In contrast, adjustment for tumor mass provided no evidence of comparable protection in adipose tissue. These findings indicate that the cachectic host does not lose physiological plasticity uniformly but instead retains compartment-specific responsiveness to exercise.

UN-KC-6141 tumor progression was associated not only with skeletal muscle atrophy, but also with adipose tissue wasting, hepatic transcriptional remodeling, splenomegaly, elevated circulating IL-6 and alterations in feeding and spontaneous behavior. This integrated phenotype therefore reproduces several major components of the systemic cachectic syndrome. Notably, this phenotype developed without any reduction in food intake, indicating that wasting was not secondary to anorexia.

A central finding is that skeletal muscle remained responsive to voluntary exercise despite ongoing tumor progression and systemic cachectic stress. Exercise has been proposed as a strategy to counteract cancer-associated muscle wasting through effects on muscle protein turnover, inflammatory signaling, oxidative metabolism and mitochondrial function, although preclinical evidence remains heterogeneous across tumor models and exercise protocols (Niels *et al*., 2020; Collao *et al*., 2023; Tamayo-Torres *et al*., 2024). More recently, Tsitkanou et al. (2026) reported that wheel-based exercise initiated before and maintained during tumor progression preserved muscle mass and reduced tumor burden in C26-bearing mice. The slower tumor growth observed with VWR is consistent with previous evidence that voluntary running can suppress tumor growth, notably through epinephrine- and IL-6-dependent mobilization of natural killer cells (Pedersen *et al*., 2016), although tumor immunity was not assessed here. Because VWR also reduced tumor burden, its effect on muscle could have reflected reduced disease severity, but gastrocnemius mass remained greater after adjustment for tumor mass. Because low-field NMR includes the tumor in the lean compartment, the conclusions regarding skeletal muscle rely on directly weighed muscles.

The molecular profile of skeletal muscle was consistent with this preserved responsiveness. Activation of proteolytic and autophagy-related pathways contributes to skeletal muscle remodeling during cancer cachexia, with the ubiquitin ligases *Fbxo32*/atrogin-1 and *Trim63*/MuRF1 frequently associated with experimental muscle wasting (Baracos *et al*., 2018; Tamayo-Torres *et al*., 2024). In the present study, VWR was associated with lower *Fbxo32*, *Trim63* and *Map1lc3b* gene expression, whereas AKT phosphorylation was not clearly altered. These findings are compatible with reduced expression of genes associated with muscle catabolic remodeling rather than major activation of canonical AKT-dependent anabolic signaling. The lower Eif4ebp1 expression should be interpreted cautiously, as it does not directly indicate translational activity. Interestingly, direct comparison of VWR and TR in healthy mice has also revealed paradigm-specific skeletal muscle adaptations, including reductions in *Fbxo32* expression in the tibialis anterior following VWR that were not reproduced by TR exercise (Schmitt *et al*., 2020). Thus, the molecular divergence observed here may reflect, at least in part, differences in the muscular stimulus generated by the two exercise paradigms, although lower Trim63 expression was also observed in CaTR mice.

Importantly, preservation of skeletal muscle mass was not accompanied by recovery of forelimb grip strength. Muscle function depends not only on muscle mass but also on muscle quality, neural activation, fatigue, motivation and the broader systemic environment (Martin & Freyssenet, 2021), all of which may constrain functional recovery during systemic disease. The present response should therefore be interpreted as preservation of skeletal muscle mass rather than restoration of muscle function.

The response of adipose tissue contrasted markedly with that of skeletal muscle. Adipose tissue wasting persisted in exercised mice, and ANCOVA provided no evidence that VWR preserved epididymal adipose tissue after accounting for tumor burden. Muscle and adipose tissue wasting in pancreatic cancer are increasingly recognized as interconnected but partially dissociable processes, adipose tissue loss occurring early in pancreatic cancer (Danai *et al*., 2018). Experimental work has demonstrated inter-organ communication between these compartments, with tumor-derived IL-6 promoting adipose lipolysis and muscle dysfunction (Rupert *et al*., 2021). Our findings extend this concept by showing that these tissues also differ in their responsiveness to exercise. In fact, after adjustment for tumor mass, epididymal adipose tissue mass was even lower in CaVWR than in CaCTL mice, suggesting that the additional energy demand of voluntary running may have further mobilized adipose lipid stores in an already catabolic context. A similar pattern was observed in CaTR mice after adjustment for tumor mass (*P* = 0.040), suggesting a contribution of exercise-related energy expenditure common to both paradigms. Because UCP1 was lower rather than higher in CaVWR mice, adipose tissue browning is unlikely to account for adipose tissue loss in this group.

A similar pattern was observed in the liver. Tumor-bearing mice displayed marked hepatic transcriptional remodeling, including alterations in acute-phase response and metabolic genes, together with a non-significant increase in hepatic cardiolipin content. Neither VWR nor TR substantially restored these changes. The liver is increasingly recognized as an active contributor to whole-body metabolic dysregulation during cancer cachexia through altered substrate metabolism, acute-phase protein synthesis, inflammatory signaling and inter-organ communication (Baracos *et al*., 2018; Gonçalves *et al*., 2024). The persistence of these hepatic abnormalities despite skeletal muscle preservation therefore indicates that VWR did not globally normalize host metabolism.

The persistence of splenomegaly and elevated circulating IL-6 further reinforces this interpretation. Yet VWR preserved skeletal muscle without a detectable reduction in circulating IL-6 or splenomegaly, whereas TR modestly reduced spleen mass without preserving skeletal muscle. Muscle preservation therefore cannot readily be attributed to normalization of the systemic inflammatory environment. Candidate local or systemic mechanisms not captured by circulating IL-6 may include myokine signaling, extracellular vesicles, neuromuscular adaptations or local stress-response pathways, although these possibilities were not directly investigated here.

These tissue-specific responses may be reconciled by considering the nature of the signal that each compartment receives during exercise. Skeletal muscle is the only compartment exposed to a direct, local contractile and mechanical stimulus, which may be sufficient to counteract catabolic signaling even within a persistently inflammatory systemic environment, as suggested by the lower expression of Fbxo32 and Trim63 in CaVWR mice without any reduction in circulating IL-6. In contrast, adipose tissue mainly experiences exercise as an additional energy demand. In a host already in negative energy balance, this demand is likely to be met, at least in part, by further mobilization of lipid stores, consistent with the lower adipose tissue mass observed at a given tumor burden in both exercised groups. CaVWR mice increased their food intake by approximately 13%, but this compensation was apparently insufficient to prevent further mobilization of lipid stores. In this view, each compartment responds according to the dominant signal it receives: a local signal for skeletal muscle and systemic signals for adipose tissue and liver. This hypothesis remains to be tested directly, for instance by dissociating local muscle loading from whole-body energy expenditure.

The behavioral phenotype further distinguished the two paradigms, possibly reflecting their different temporal organization. CaVWR mice ran predominantly during the dark phase, in phase with their natural activity rhythm, and displayed more frequent dark-phase feeding events. In contrast, CaTR mice were trained during the light (rest) phase and displayed greater light-phase food intake, without an increase in total daily intake, and greater estimated dark-phase sleep, a pattern observed on both training and non-training days, suggesting a sustained adaptation of feeding and activity rhythms to the training schedule rather than an acute post-exercise response. Their lower light-phase resting energy expenditure may indicate energy-sparing adaptations to the energy cost of training. Because CaTR mice were only about 3% lighter than CaCTL mice during metabolic phenotyping, this difference is unlikely to be explained by body mass. In addition, among tumor-bearing mice, longer dark-phase sleep was associated with greater body-weight and adipose tissue loss independently of group and tumor mass, suggesting that reduced nocturnal activity may represent a behavioral marker of cachexia severity. However, metabolic phenotyping involved only five to six mice per group; these observations should therefore be considered exploratory and do not establish a behavioral mechanism for skeletal muscle preservation. Cachexia-related behavioral regulation may involve integrated inflammatory and neuroendocrine pathways. GDF15–GFRAL signaling represents one potential pathway linking peripheral metabolic stress to appetite, body weight and sickness-related behavior (Suriben *et al*., 2020; Groarke *et al*., 2024); however, GDF15 was not measured in the present study and its potential contribution remains speculative.

The contrasting effects of VWR and TR should therefore be interpreted within the broader context of exercise paradigm- and tumor-dependent responses. In the present study, the TR protocol not only failed to reproduce the tumor and skeletal muscle adaptations associated with VWR, but was also associated with a less favorable body-weight trajectory, lower tumor-free body weight and greater fat mass loss than in inactive tumor-bearing mice. In an already catabolic host, imposed exercise performed during the rest phase may therefore carry an energetic and possibly stress-related cost that is offset neither by a local muscular benefit nor by a compensatory increase in food intake, consistent with the behavioral and metabolic profile of CaTR mice. These data nevertheless relate to one specific treadmill protocol and do not demonstrate that TR is intrinsically ineffective or detrimental in cancer cachexia. An opposite pattern has been reported in prostate cancer, where structured TR reduced tumor growth while preserving adipose tissue and muscle strength in tumor-bearing mice (Dieumegard *et al*., 2025). VWR also showed limited effects on prostate tumor responses to radiotherapy (Dufresne *et al*., 2021). In C26-bearing mice, VWR has been reported to preserve muscle mass and mitochondrial integrity (Pigna *et al*., 2016; Kitaoka *et al*., 2021). Taken together, these observations indicate that biological responses to exercise cannot be predicted from the nominal exercise paradigm alone.

The two paradigms were not matched for exercise dose. Within the CaVWR group, running distance was not associated with gastrocnemius mass or tumor-free body weight, suggesting that, beyond a certain level, additional running volume conferred no further benefit. However, most CaVWR mice ran well above the theoretical distance of a TR session (approximately 640 m), so a threshold effect of exercise volume cannot be excluded as a contributor to the divergent effects of the two paradigms. Because the two paradigms also differed in several other characteristics (see Limitations), these findings should not be interpreted as evidence that voluntary exercise is intrinsically superior to treadmill exercise. Rather, they emphasize that exercise prescription itself should be considered a biological variable in preclinical exercise oncology.

### Limitations

Several limitations should be acknowledged. First, the study was performed in a single syngeneic pancreatic cancer model and exclusively in male mice, limiting generalization across tumor models and sexes. Second, although ANCOVA strengthens the interpretation that skeletal muscle mass preservation was not solely attributable to reduced tumor burden, statistical adjustment cannot establish causal independence between tumor control and muscle preservation. Third, VWR and TR were not matched for exercise volume, intensity, temporal distribution or circadian timing, preventing identification of the specific exercise characteristic responsible for their divergent responses, although the absence of a dose–response relationship within the CaVWR group argues against exercise volume as the sole determinant. Fourth, the molecular analyses provide information on selected markers but do not directly measure muscle protein synthesis and proteolytic or autophagic fluxes, and therefore cannot establish the mechanisms responsible for muscle mass preservation. Fifth, the mechanisms underlying the tissue-specific responsiveness observed across skeletal muscle, adipose tissue and liver were not directly investigated. Sixth, tumors were implanted subcutaneously rather than orthotopically, which does not reproduce the pancreatic tumor microenvironment, and no exercising healthy groups were included. Finally, sleep was estimated from metabolic-cage behavioral monitoring rather than measured using electrophysiological recordings, body composition was assessed by low-field NMR, which includes the subcutaneous tumor in lean and fluid compartments, metabolic phenotyping was restricted to five to six mice per group, and food intake was estimated by weighing the food provided, which may overestimate intake in case of food spillage. Future studies should determine the local and systemic signals governing tissue-specific exercise responsiveness during cancer cachexia and whether the retained plasticity of skeletal muscle can be harnessed to improve both tissue preservation and functional outcomes.

## CONCLUSION

VWR does not globally reverse pancreatic cancer cachexia but selectively preserves skeletal muscle within a persistently altered systemic environment. This muscle preservation remains evident after adjustment for tumor burden, whereas adipose tissue wasting is not prevented, and is even accentuated at a given tumor burden, and hepatic metabolic remodeling and systemic inflammation are largely unaffected. These findings reveal marked tissue-specific differences in exercise responsiveness during pancreatic cancer cachexia and indicate that skeletal muscle retains substantial physiological plasticity despite ongoing systemic disease. More broadly, they emphasize that both the physiological compartment examined and the characteristics of the exercise paradigm are important determinants of tumor–host adaptation. From a clinical perspective, the accentuated loss of adipose tissue observed with exercise suggests that exercise interventions in patients with pancreatic cancer cachexia should be combined with adequate nutritional support, consistent with current multimodal approaches to cachexia management.

## ACKNOWLEDGMENTS

The authors thank Pascale Bellaud (H2P2 histopathology core facility, BIOSIT, Université de Rennes, Rennes, France) and Florence Boutillon (ARCHE animal facility, BIOSIT, Université de Rennes, Rennes, France) for their technical help. We also thank Julie Massart (Institut NUMECAN, Université de Rennes, Rennes, France) for discussion. The authors thank Prof. S.K. Batra for providing the UN-KC-6141 cell line (MTA, University of Nebraska Medical Center, 986099 Nebraska Medical Center, Omaha, NE 68198-6099, USA). This work was funded by Inserm (Institut national de la santé et de la recherche médicale), the Ligue contre le cancer (Ille-et-Vilaine and Sarthe departmental committees) and Institut Universitaire de France.

## AUTHOR CONTRIBUTIONS

Conception or design of the work: A.R., C.R., P.L., A.C. and C.Ri. Acquisition, analysis or interpretation of data for the work: C.R., A.D., L.-A.C., L.L.-O., B.M., L.R., S.S., M.V., G.A. and A.R. Drafting the work: C.R., A.D. and A.R. Revising it critically for important intellectual content: all authors. All authors have read and approved the final version of this manuscript and agree to be accountable for all aspects of the work in ensuring that questions related to the accuracy or integrity of any part of the work are appropriately investigated and resolved. All persons designated as authors qualify for authorship, and all those who qualify for authorship are listed.

## DATA AVAILABILITY STATEMENT

The data that support the findings of this study are available from the corresponding author upon reasonable request.

## REFERENCES

American Cancer Society (2026). Cancer Facts & Figures. Available at: https://www.cancer.org/content/dam/cancer-org/research/cancer-facts-and-statistics/annual-cancer-facts-and-figures/2026/2026-cancer-facts-and-figures.pdf.

Assi M, Derbré F, Lefeuvre-Orfila L & Rébillard A (2016). Antioxidant supplementation accelerates cachexia development by promoting tumor growth in C26 tumor-bearing mice. Free Radic Biol Med 91, 204–214.

Baracos VE, Martin L, Korc M, Guttridge DC & Fearon KCH (2018). Cancer-associated cachexia. Nat Rev Dis Primer 4, 17105.

Bligh EG & Dyer WJ (1959). A rapid method of total lipid extraction and purification. Can J Biochem Physiol 37, 911–917.

Collao N, Sanders O, Caminiti T, Messeiller L & De Lisio M (2023). Resistance and endurance exercise training improves muscle mass and the inflammatory/fibrotic transcriptome in a rhabdomyosarcoma model. J Cachexia Sarcopenia Muscle 14, 781– 793.

Danai LV et al. (2018). Altered exocrine function can drive adipose wasting in early pancreatic cancer. Nature 558, 600–604.

Dieumegard A, Dufresne S, Richard C, Orfila L, Martin B, Rouvière L, Oliveira AR, Croyal M, Mathieu R & Rébillard A (2025). Structured Treadmill Training as a Strategy to Mitigate Tumor Growth and Preserve Adipose tissue and Muscle Strength in Prostate Tumor Bearing Mice. Med Sci Sports Exerc; DOI: 10.1249/MSS.0000000000003675.

Dufresne S, Richard C, Dieumegard A, Orfila L, Delpon G, Chiavassa S, Martin B, Rouvière L, Escoffre J-M, Oujagir E, Denis de Senneville B, Bouakaz A, Rioux-Leclercq N, Potiron V & Rébillard A (2021). Voluntary Wheel Running Does Not Enhance Radiotherapy Efficiency in a Preclinical Model of Prostate Cancer: The Importance of Physical Activity Modalities? Cancers 13, 5402.

Fearon K, Strasser F, Anker SD, Bosaeus I, Bruera E, Fainsinger RL, Jatoi A, Loprinzi C, MacDonald N, Mantovani G, Davis M, Muscaritoli M, Ottery F, Radbruch L, Ravasco P, Walsh D, Wilcock A, Kaasa S & Baracos VE (2011). Definition and classification of cancer cachexia: an international consensus. Lancet Oncol 12, 489–495.

Gonçalves DC, Gomes SP & Seelaender M (2024). Metabolic, Inflammatory, and Molecular Impact of Cancer Cachexia on the Liver. Int J Mol Sci 25, 11945.

Groarke JD, Crawford J, Collins SM, Lubaczewski S, Roeland EJ, Naito T, Hendifar AE, Fallon M, Takayama K, Asmis T, Dunne RF, Karahanoglu I, Northcott CA, Harrington MA, Rossulek M, Qiu R & Saxena AR (2024). Ponsegromab for the Treatment of Cancer Cachexia. N Engl J Med 391, 2291–2303.

von Haehling S, Anker MS & Anker SD (2016). Prevalence and clinical impact of cachexia in chronic illness in Europe, USA, and Japan: facts and numbers update 2016. J Cachexia Sarcopenia Muscle 7, 507–509.

Kitaoka Y, Miyazaki M & Kikuchi S (2021). Voluntary exercise prevents abnormal muscle mitochondrial morphology in cancer cachexia mice. Physiol Rep 9, e15016.

Martin A & Freyssenet D (2021). Phenotypic features of cancer cachexia-related loss of skeletal muscle mass and function: lessons from human and animal studies. J Cachexia Sarcopenia Muscle 12, 252–273.

Niels T, Tomanek A, Freitag N & Schumann M (2020). Can Exercise Counteract Cancer Cachexia? A Systematic Literature Review and Meta-Analysis. Integr Cancer Ther 19, 1534735420940414.

Pedersen L, Idorn M, Olofsson GH, Lauenborg B, Nookaew I, Hansen RH, Johannesen HH, Becker JC, Pedersen KS, Dethlefsen C, Nielsen J, Gehl J, Pedersen BK, thor Straten P & Hojman P (2016). Voluntary Running Suppresses Tumor Growth through Epinephrine- and IL-6-Dependent NK Cell Mobilization and Redistribution. Cell Metab 23, 554–562.

Pigna E, Berardi E, Aulino P, Rizzuto E, Zampieri S, Carraro U, Kern H, Merigliano S, Gruppo M, Mericskay M, Li Z, Rocchi M, Barone R, Macaluso F, Di Felice V, Adamo S, Coletti D & Moresi V (2016). Aerobic Exercise and Pharmacological Treatments Counteract Cachexia by Modulating Autophagy in Colon Cancer. Sci Rep 6, 1–14.

Rupert JE, Narasimhan A, Jengelley DHA, Jiang Y, Liu J, Au E, Silverman LM, Sandusky G, Bonetto A, Cao S, Lu X, O’Connell TM, Liu Y, Koniaris LG & Zimmers TA (2021). Tumor-derived IL-6 and trans-signaling among tumor, fat, and muscle mediate pancreatic cancer cachexia. J Exp Med 218, e20190450.

Schmitt A, Herzog P, Röchner F, Brändle A-L, Fragasso A & Munz B (2020). Skeletal muscle effects of two different 10-week exercise regimens, voluntary wheel running, and forced treadmill running, in mice: A pilot study. Physiol Rep 8, e14609.

Suriben R et al. (2020). Antibody-mediated inhibition of GDF15-GFRAL activity reverses cancer cachexia in mice. Nat Med 26, 1264–1270.

Tamayo-Torres E, Garrido A, de Cabo R, Carretero J & Gómez-Cabrera MC (2024). Molecular mechanisms of cancer cachexia. Role of exercise training. Mol Aspects Med 99, 101293.

Torres MP, Rachagani S, Souchek JJ, Mallya K, Johansson SL & Batra SK (2013). Novel pancreatic cancer cell lines derived from genetically engineered mouse models of spontaneous pancreatic adenocarcinoma: applications in diagnosis and therapy. PloS One 8, e80580.

Tsitkanou S, Koopmans PJ, Cabrera AR, Serrano N, Peterson CS, Muhyudin R, Malik ZB, Goeke M, Schrems ER, Morena F, Wen Y, Washington TA, Murach KA & Greene NP (2026). Exercise training prior to and during cancer in mice preserves muscle mass, reduces tumour weight and suppresses molecular mediators of cachexia. J Physiol 604, 6028–6056.

