## Supplementary material for "Voluntary wheel running selectively preserves skeletal muscle during pancreatic cancer cachexia despite persistent systemic alterations": Figure S1

**Supplemental data**

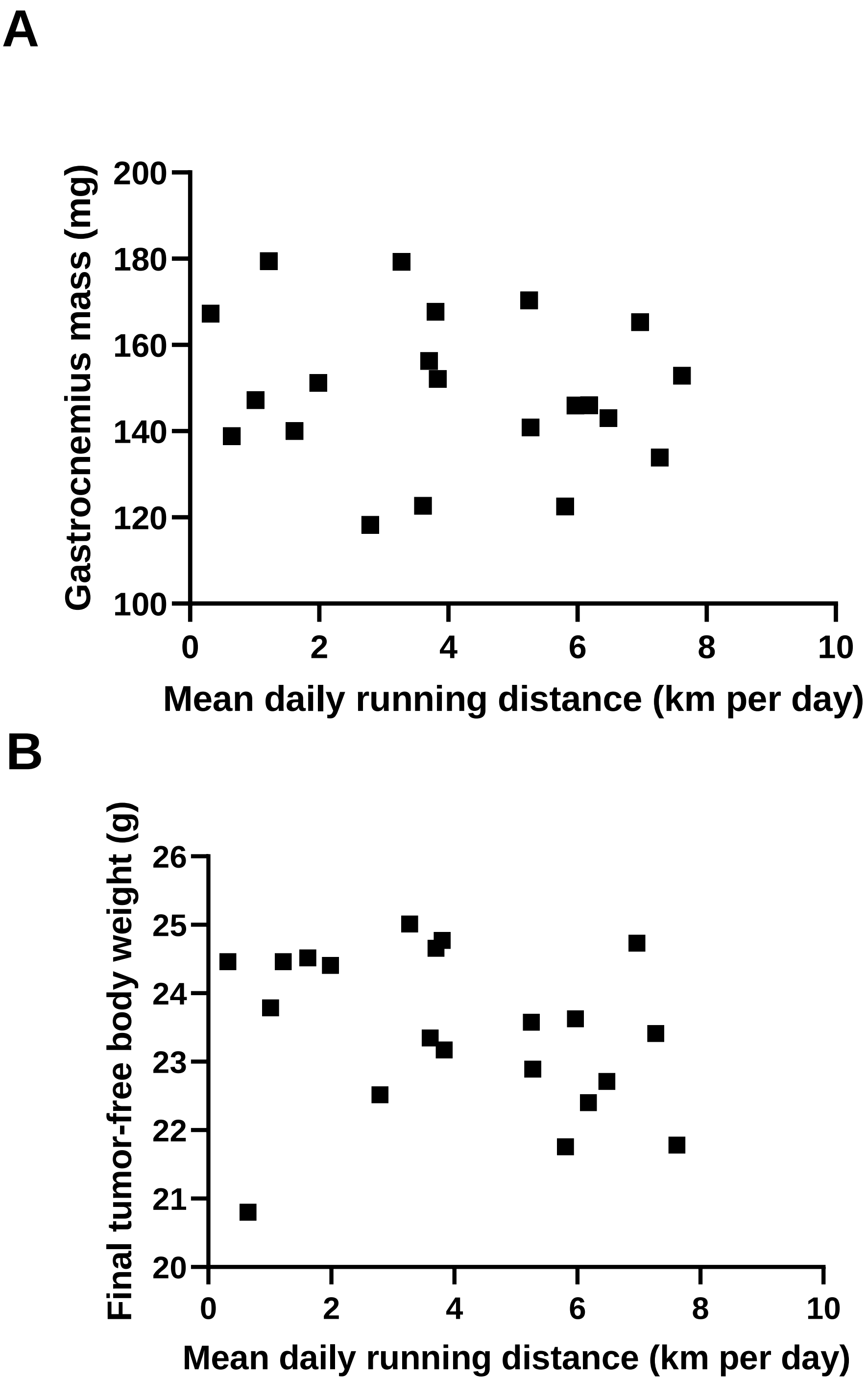


**Figure S1. Voluntary running distance, gastrocnemius mass and tumor-free body weight in CaVWR mice.** Relationship between mean daily running distance and (A) gastrocnemius mass and (B) final tumor-free body weight (n = 21; Pearson correlation). Running distance was calculated from daily wheel revolutions and averaged over the intervention period.
